# *Gardnerella* vaginolysin potentiates glycan molecular mimicry by *Neisseria gonorrhoeae*

**DOI:** 10.1101/2023.04.15.537036

**Authors:** Sydney Morrill, Sudeshna Saha, Ajit P Varki, Warren G. Lewis, Sanjay Ram, Amanda L. Lewis

## Abstract

Bacterial vaginosis (BV) is a condition of the vaginal microbiome in which there are lower levels of “healthy” *Lactobacillus* species and an outgrowth of diverse anaerobic bacteria. BV is associated with increased risk of infection by the bacterium *Neisseria gonorrhoeae* - the causative agent of gonorrhea. Here we test if one known facet of BV - the presence of bacterial cytolysins – leads to the mobilization of specific intracellular contents that aid in gonococcal virulence. We cloned and expressed recombinant vaginolysin (VLY), a cytolysin produced by the BV-associated bacterium *Gardnerella*, verifying that it liberates the contents of red blood cells and cervical epithelial (HeLa) cells while vector control preparations made in parallel did not. We tested if VLY mediates a well-known virulence mechanism of gonococcus – the molecular mimicry of host glycans. To evade host immunity, *N. gonorrhoeae* caps its surface lipooligosaccharide (LOS) with α2-3-linked sialic acid. To do this, gonococci must scavenge an intermediate metabolite made and used inside host cells. Flow-cytometry based lectin-binding assays showed that, compared to controls, gonococci exposed to vaginolysin-liberated contents of HeLa cells displayed greater sialic acid capping of their LOS. This higher level of bacterial sialylation was accompanied by increased binding of the complement regulatory protein Factor H, and greater resistance to complement attack. Together these results suggest that cytolytic activities present during BV may enhance the ability of *N. gonorrhoeae* to capture intracellular metabolites and evade host immunity via glycan molecular mimicry.

## Introduction

In most humans the vaginal microbiome is dominated by *Lactobacillus* species [1]. High levels of *Lactobacillus* are associated with optimal sexual and reproductive health outcomes in a wide variety of settings globally [2]. In contrast, nearly 1/3 of reproductive age women in the U.S. are affected by bacterial vaginosis (BV), a condition of vaginal microbiota [3]. In BV there is a lower relative abundance of *Lactobacillus*, especially *L. crispatus*. and outgrowth of diverse anaerobic bacteria [1, 4]. This shift in microbial composition is often accompanied by higher vaginal pH (>4.5), a fishy odor, changes in vaginal mucus characteristics and epithelial phenotypes [4, 5]. BV has been associated with numerous adverse health outcomes, including increased risk of infection by the bacterium *Neisseria gonorrhoeae* [6].

*Neisseria gonorrhoeae* is the causative agent of the uniquely human gonorrhea, a prolific STI with upwards of 80 million new cases per year worldwide [7]. Some individuals with gonorrhea infection of the cervicovaginal tract can experience severe consequences, including pelvic inflammatory disease [8], ectopic pregnancy [9] and in rare cases, bacteremia [10]. There is currently no commercially available vaccine for *N. gonorrhoeae* infection, and resistant strains have been found against almost every available antibiotic [7]. There have been numerous clinical studies indicating that BV is a risk factor for co-incident and new gonorrhea infection, even after adjusting for relevant confounders [6, 11-13]. However, the mechanisms by which BV predisposes individuals to gonococcal infection have not been well defined.

Some of the abundant bacteria in BV produce cytolytic proteins. In particular, vaginolysin (VLY) is a cytolysin produced by members of the genus *Gardnerella*, which are often identified as the most abundant members of the BV consortium [1, 14]. In other biological contexts, cytolytic proteins can aid bacterial colonization or virulence, for example by lysing host red blood cells (RBCs) and promoting iron-acquisition by liberating intracellular heme [15]. We hypothesized that cytolytic proteins in BV may be aiding in gonococcal colonization of the female reproductive tract by disrupting genital epithelial cells and freeing host intracellular contents that may aid in acquisition of virulence traits. To test this hypothesis mechanistically, we specifically investigated if *Gardnerella* VLY enhances resistance of *N. gonorrhoeae* to attack by the host complement system using a well-described mechanism of immune evasion in which the bacterium scavenges carbohydrate building blocks from host cells in a ploy to decorate its own surface with host-like glycans.

*N. gonorrhoeae* engages in a feat of molecular mimicry by directly transferring sialic acid residues to its surface lipooligosaccharide (LOS) [16]. *N*-acetylneuraminic acid or Neu5Ac is the major sialic acid found in humans. It is present on all cell surfaces and particularly abundant at mucosal surfaces, including the genital tract. Factor H is found in mammalian body fluids and inhibits the host complement system by recognizing sialic acids and preventing the complement system from targeting “self” cells as foreign [17]. *N. gonorrhoeae* mimics the appearance of host glycans by scavenging a metabolically activated form of sialic acid, cytidine-5’-monophospho-N-acetyl neuraminic acid (CMP-Neu5Ac) to cap its LOS with Neu5Ac [16]. *N. gonorrhoeae* lacks the ability to synthesize CMP-Neu5Ac, and therefore must scavenge this molecule from host cells [18].

In this study we use an *in vitro* model to test if VLY treatment of cervical epithelial cells potentiates *N. gonorrhoeae* survival by enabling resistance to the complement system via glycan molecular mimicry. If VLY treatment of host cells is able to enhance CMP-Neu5Ac capture and LOS sialylation, we expect to see: 1) capping of surface galactose residues, 2) higher α2-3 surface sialylation that is dependent on its surface sialyltransferase (Lst), 3) more binding to Factor H, and 4) greater resistance to complement attack.

## Results

To test our working hypothesis, that *Gardnerella* VLY potentiates a well-known mechanism of gonococcal molecular mimicry, we first exposed human red blood cells (hRBCs) and cervical epithelial cells to purified recombinant VLY expressed in *E. coli* and measured the release of intracellular contents. As described in the methods, we began by cloning and optimizing expression of VLY (sequence shown in Supplementary Figure 1). To avoid biological effects that might be elicited by endotoxin, a modified *E. coli* strain containing a detoxified form of lipopolysaccharide was used [19]. As an additional control, the empty plasmid vector was transformed into the same *E. coli* expression strain and identical parallel purification procedures were performed. Preparations from cells containing the VLY construct revealed a single band at the expected molecular weight (about 53 kD) in SDS-PAGE, while the vector-only control preparation yielded no bands (Fig. 1A). As expected, incubating recombinant VLY with hRBCs led to heme release (Figure 1B). Maximum heme release from hRBCs was achieved between 50 - 100 ng/ml VLY, with an EC50 of 4.68 ng/ml (95% CI 4.18 - 5.26 ng/ml, hillslope 1.95).

**Figure 1:**
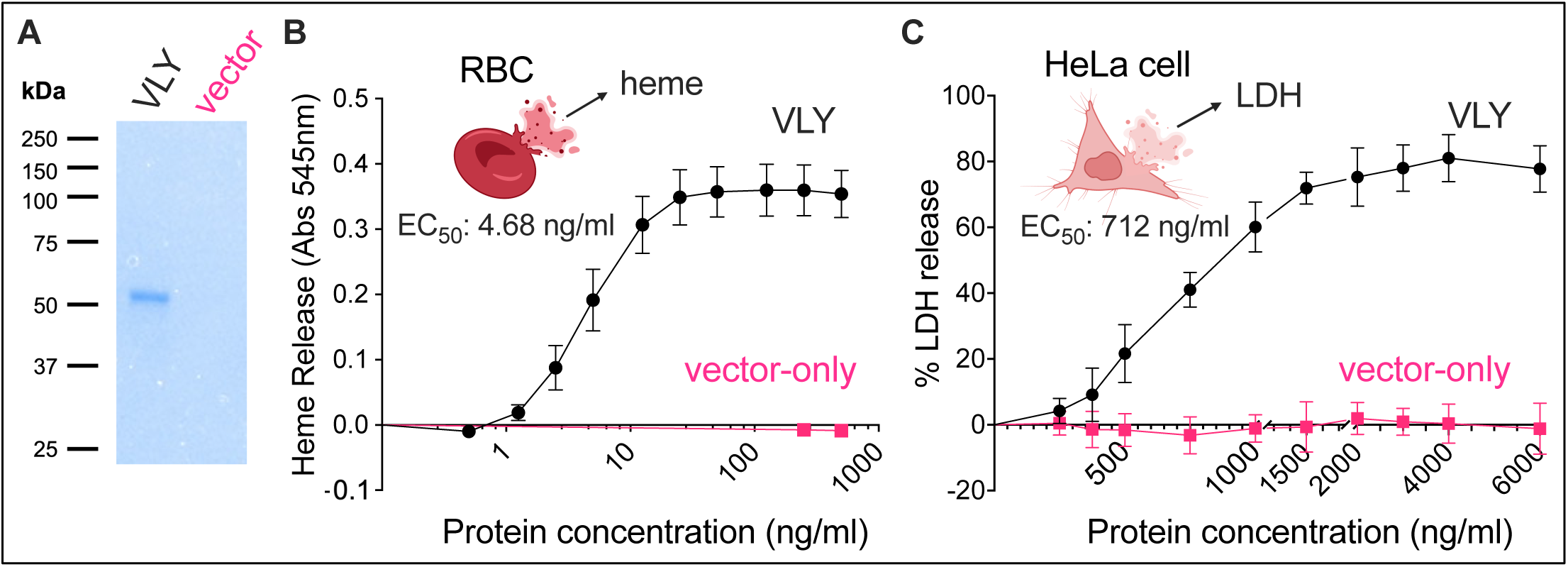
Exposure of human cells to *Gardnerella* vaginolysin leads to a release of intracellular contents from human red blood cells and cervical epithelial cells. (A) Coomassie blue SDS-PAGE gel showing recombinant purified vaginolysin (VLY) and a parallel control preparation using cells containing the empty expression vector. (B) Cytolytic activity of VLY against human RBCs determined by heme release (detected as absorbance at 545 nm). The EC50 was calculated using a four-parameter nonlinear regression model. The data is a combination of three independent experiments, with three biological replicates per experiment. (C) Cytolytic activity of VLY against HeLa cells determined by LDH release. The EC50 was calculated using a four-parameter nonlinear regression model. Data is a combination of two independent experiments, with three to four biological replicates per experiment.

VLY was also active against HeLa cells, a human cervical epithelial cell line, as measured by the release of a cytosolic enzyme, lactate dehydrogenase (LDH), often used as a marker of cell membrane damage. Compared to hRBCs, HeLa cells required 20-30-fold more VLY (1500 – 2000 ng/ml VLY) to release intracellular contents. Likewise, the EC50 of VLY against HeLa cells (712 ng/ml) was about 150-fold higher compared to hRBCs (95% CI 670 -757 ng/ml, hillslope 3.02).

To study structural elements of the gonococcal LOS relevant to our hypothesis, we developed a lectin-binding assay in flow cytometry to monitor terminal α2-3-linked sialic acids (Fig. 2A). Prior studies have shown that at baseline, gonococcal lacto-*N*-neotetraose (LNnT) LOS terminates with galactose (Gal) in a β1-4 linkage to N-acetylglucosamine (GlcNAc). When CMP-Neu5Ac is provided or becomes available in the host, the gonococcal LOS-sialyltransferase (Lst) adds Neu5Ac to the terminal Gal in an α2-3 linkage (Fig. 2A). *Maackia amurensis* Lectin I (MAL-I) recognizes the Neu5Acα2-3Galβ1-4GlcNAc moiety that terminates sialylated LOS (Fig. 2A). This flow cytometry-based assay shows that MAL-I binds to *N. gonorrhoeae* (strain F62) provided with exogenous CMP-Neu5Ac in a dose-dependent manner (Fig. 2B). Additional experiments demonstrated that this increase in MAL-I-binding requires Lst (Supplementary Figure 2).

**Figure 2:**
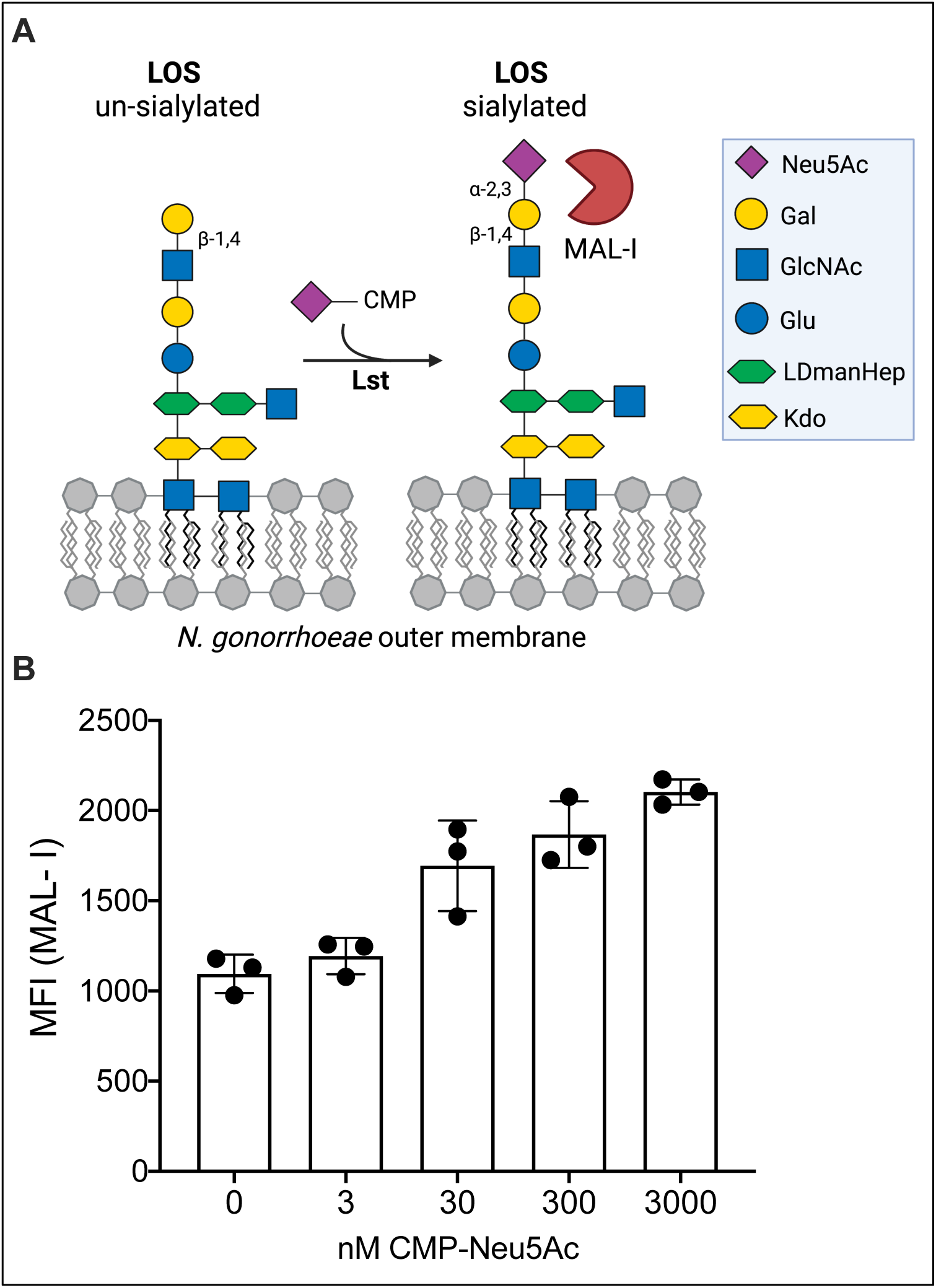
MAL-I binds to sialylated gonococcal LOS. (A) Schematic of Neu5Ac addition to the terminal lactosamine of gonococcal LOS. MAL-I binding site is indicated. (B) MAL-I binding to *N. gonorrhoeae* F62 WT following bacterial incubation with increasing concentrations of CMP-Neu5Ac. Presented as median fluorescence intensity (MFI). Data is from a single experiment, with three biological replicates per CMP-Neu5Ac concentration.

We hypothesized that exposure of epithelial cells to VLY would create conditions in which the gonococcus can more readily place sialic acids onto its LOS. To test this, *N. gonorrhoeae* was incubated with supernatants of cervical epithelial cells treated with VLY. The general method for this schematic is represented in Figure 3A. Briefly, HeLa cells were treated with 1 µg/ml VLY, or with an equivalent dilution of the control preparation from cells harboring the empty vector. After 90 minutes of incubation, HeLa supernatants were collected, centrifuged, and filtered to remove cell-debris >0.22 µm. *N. gonorrhoeae* strain F62 was then incubated with supernatants from cells treated with VLY or the vector control preparation and MAL-I binding was evaluated. Consistent with our hypothesis, gonococcus exhibited greater binding to MAL-I after exposure to VLY-extracted cellular contents compared to vector controls (Fig 3B & 3D). The VLY-mediated increase in MAL-I binding was almost entirely abrogated by deleting the gene encoding Lst (Fig. 3C-D). We also observed a small but significant source of Lst-independent MAL-I binding, which is discussed in in Conclusions.

**Figure 3:**
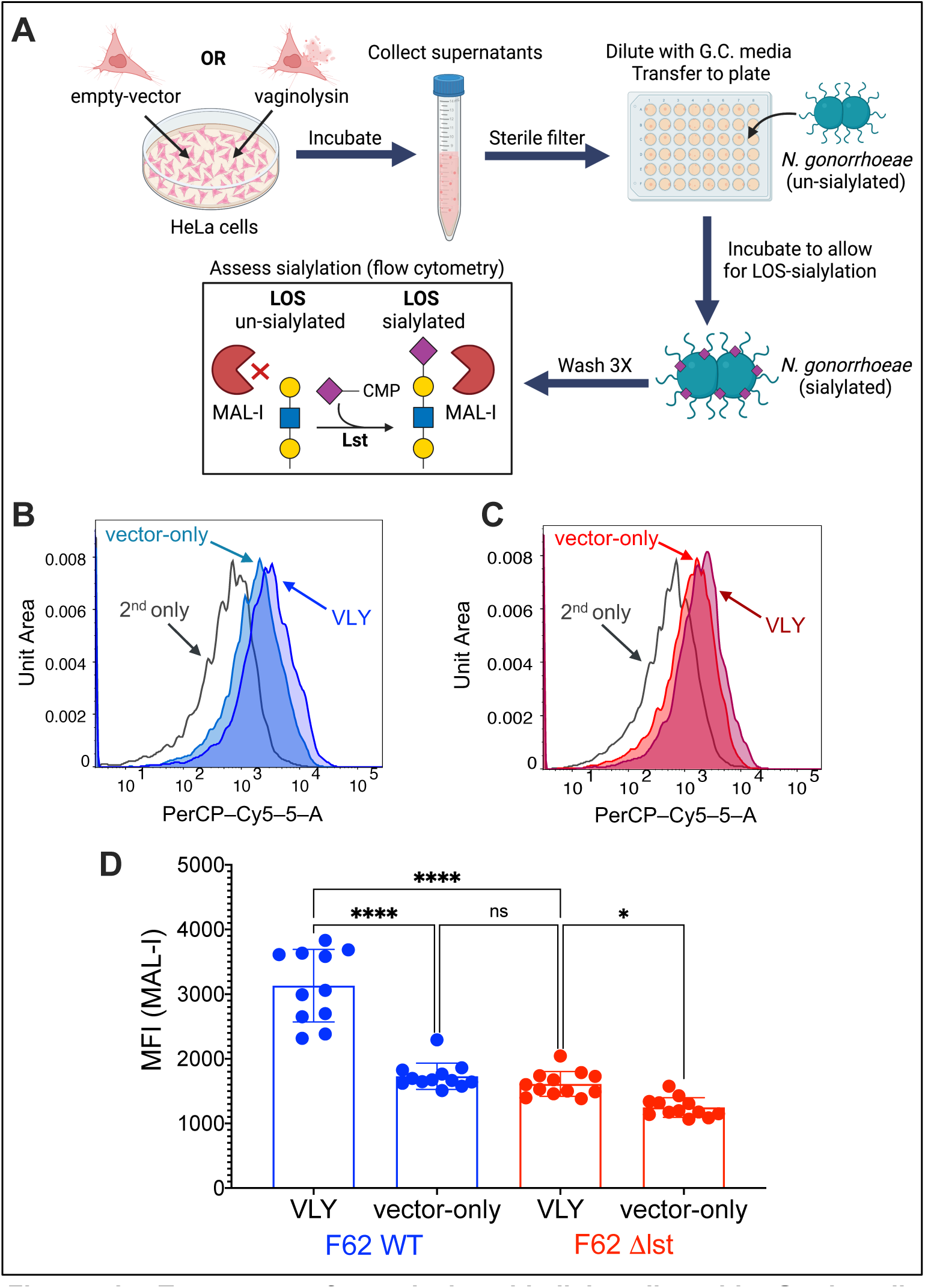
Treatment of cervical epithelial cells with Gardnerella vaginolysin potentiates Lst-dependent *N. gonorrhoeae* sialylation. N. gonorrhoeae F62 WT and Δlst were incubated with the supernatants of HeLa cells that had been exposed to either 1 μg/mL vaginolysin or a parallel dilution of the vector control preparation. (A) Schematic of the method for Figure 3 experiments, and those shown in Figure 4C and 4D. (B-D) MAL-I binding to *N. gonorrhoeae* F62 WT and Δlst after incubation with HeLa cell supernatants. (B & C) Representative histograms of MAL-I binding to (B) F62 WT or (C) Δlst. (D) MAL-I binding presented as median fluorescence intensity (MFI). Data are a combination of two independent experiments, with 3-4 biological replicates per experiment. Statistical significance was evaluated using ordinary one-way ANOVA. *p<0.05 ****p<0.0001 ns = no significance

In other experiments, *Erythrina cristagalli* lectin (ECA) was used to probe surface glycan structures of *N. gonorrhoeae*. ECA recognizes the non-sialylated LOS structure containing terminal Galβ1-4GlcNAc (Fig. 4A). As expected, ECA binding to *N. gonorrhoeae* F62 was highest in the absence of CMP-Neu5Ac, while incubation with increasing concentrations of CMP-Neu5Ac resulted in a dose dependent reduction in ECA binding as the galactose residues became capped with sialic acids (Fig. 4B). Likewise, binding of ECA to gonococci was lower after exposure to VLY-extracted cellular contents compared to vector controls (Fig. 4C-D). In both instances, the difference in ECA binding was dependent on Lst activity (Supplementary Figure 3). Together, these data suggest that the exposure of cervical epithelial cells to *Gardnerella* VLY enhances gonococcal LOS sialylation, indicated by higher MAL-I binding and lower ECA binding.

**Figure 4:**
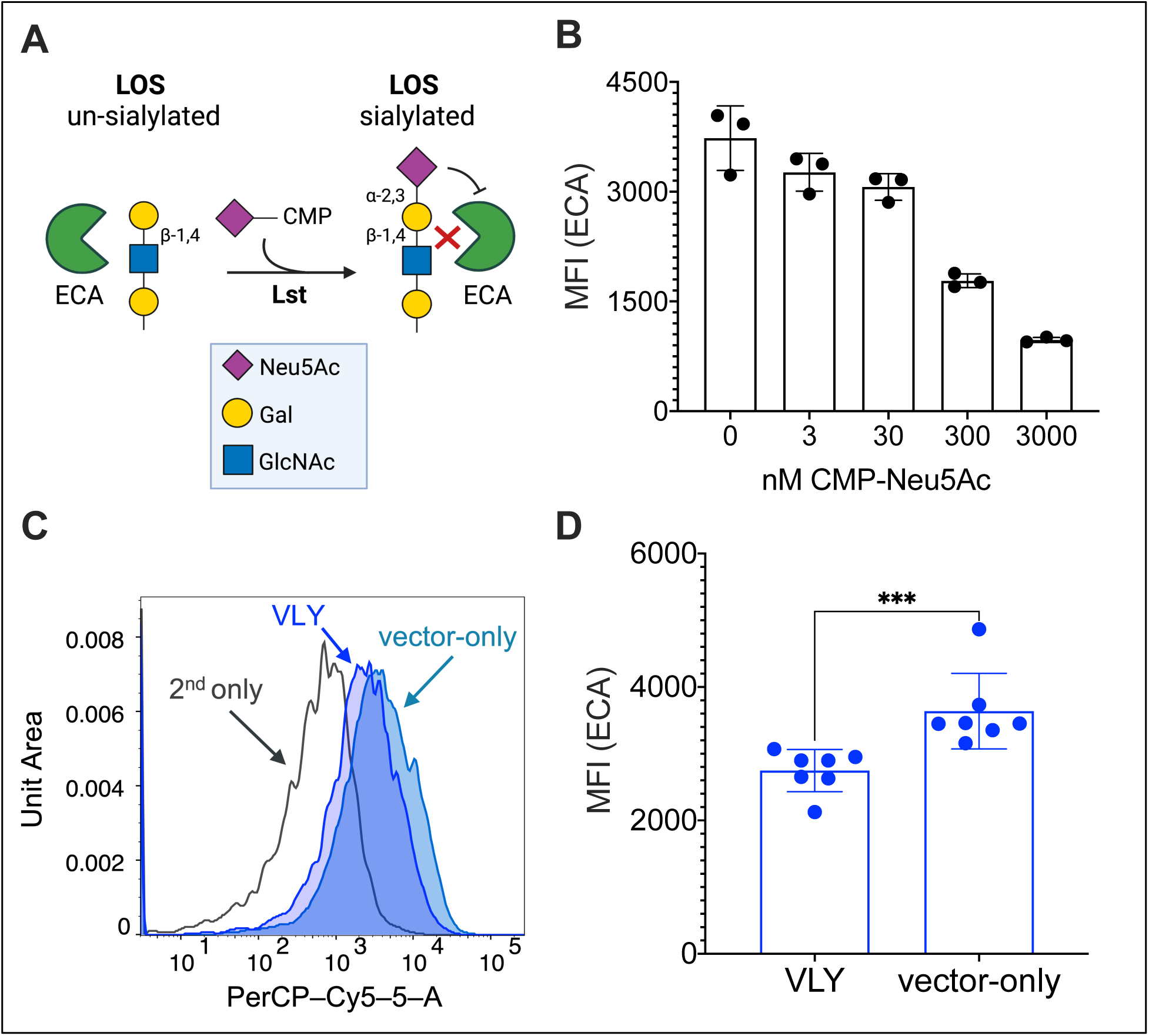
N. gonorrhoeae caps LOS galactose when exposed to VLY-liberated HeLa contents. ECA recognizes the un-sialylated terminal epitope of gonococcal LOS, and therefore binding of ECA acts as a negative readout of gonococcal sialylation. (A) Diagram of *N. gonorrhoeae* un-sialylated LOS, with ECA-binding epitope indicated. (B) ECA binding to *N. gonorrhoeae* F62 WT following bacterial incubation with increasing concentrations of CMP-Neu5Ac. Presented as median fluorescence intensity (MFI). Data is from a single experiment, with three biological replicates per concentration. (C & D) ECA binding after *N. gonorrhoeae* F62 WT was incubated with the supernatants of HeLa cells that had been exposed to either 1 μg/mL vaginolysin or a parallel dilution of the pET28a vector-only control (C) Representative histogram of ECA binding to F62 WT. (D) ECA binding presented as median fluorescence intensity (MFI). Data is a combination of two independent experiments, with 3-4 biological replicates per experiment. Significance determined by Mann-Whitney. ***p<0.001

One of the major mechanisms by which LOS sialylation potentiates gonococcal virulence is via recruitment of host Factor H (FH). The addition of α2-3-linked Neu5Ac to LNnT LOS increases binding of FH to the gonococcal surface [20], where it inhibits activation of the alternative complement pathway and protects the gonococcus from killing by the complement cascade [17]. We hypothesized that the enhanced gonococcal sialylation observed in Figures 3 and 4 upon exposure to VLY-liberated HeLa cell contents would also lead increased recruitment of FH. To test this hypothesis we used a chimeric FH protein (Fc3/FH*or S2534) [21] comprised of the Fc region of human IgG3 fused to domains 18-20 of human Factor H, which are known to interact sialylated gonococcal surface (Fig. 5A) [22]. Because S2534 is being developed as an anti-gonococcal immunotherapeutic, a point mutation (D-to-G at position 1119 in domain 19) was introduced to abrogate binding to host cells without affecting binding to sialylated gonococci. Prior studies have shown that Fc3/FH* binds to *N. gonorrhoeae* in a sialylation-dependent manner [21], which we confirmed in our assay system (Fig. 5B).

**Figure 5:**
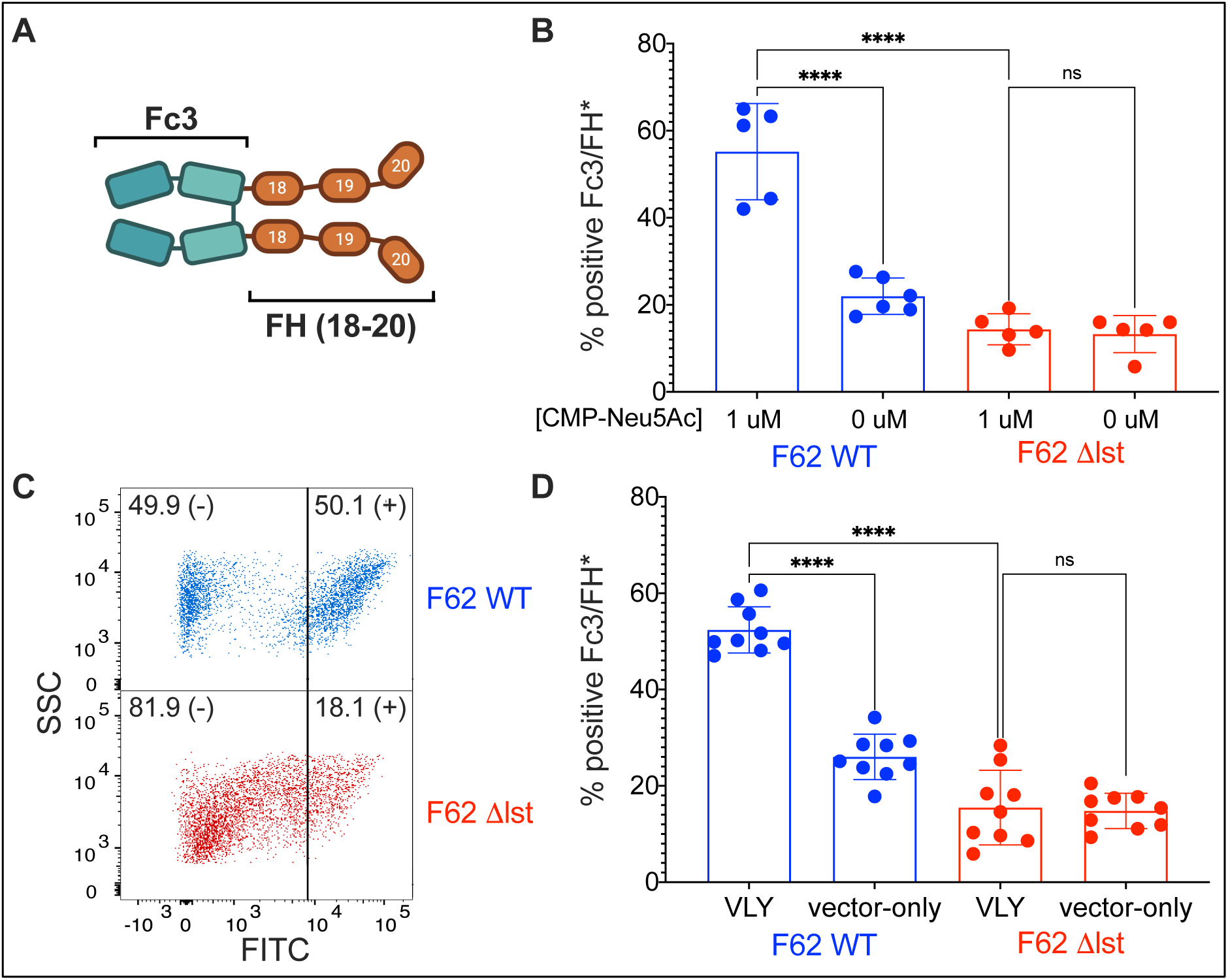
Exposure of N. gonorrhoeae to VLY-liberated HeLa contents increases sialylation-dependent recruitment of human factor H. (A) Schematic of chimeric FH protein S2534 (Fc3/FH*). S2534 consists of human IgG3 Fc receptor (Fc3) fused to the three C-terminal domains 18-20 of human factor H (FH 18-20). A point mutation in FH domain 19 (D-to-G at position 1119), which was introduced to abrogate binding to host cells, does not interfere with binding of the C-terminus of FH to sialylated *N. gonorrhoeae*. (B) Binding of Fc3/FH* to *N. gonorrhoeae* F62 WT and Δlst following incubation with either 0 μM or 1 μM CMP-Neu5Ac. Data is presented as the percentage of gated events positive for FITC. (C & D) Fc3/FH* binding after *N. gonorrhoeae* F62 WT and Δlst were incubated with the supernatants of HeLa cells that had been exposed to either 1 μg/mL vaginolysin or a parallel dilution of the pET28a vector-only control (C) Representative dot-plot of Fc3/FH* binding to *N. gonorrhoeae* WT (blue) and Δlst (red) after exposure to VLY-extracted HeLa contents. (D) Fc3/FH* binding presented as the percentage of gated events positive for FITC. (B & D) Data are a combination of two independent experiments, with 2-3 (B) or 4-5 (D) biological replicates per experiment. Significance determined by one-way ANOVA. ****p<0.0001 ns = no significance

Using the same method depicted in Figure 2A, we assessed recruitment of chimeric Fc3/FH* to *N. gonorrhoeae* F62 following bacterial exposure to VLY-liberated HeLa contents. As predicted, Fc3/FH* binding was greatly increased after exposure to the VLY-extracted HeLa contents compared to the vector control, and this effect was Lst-dependent (Fig. 5C-D).

Finally, we assessed how exposure to VLY-liberated HeLa contents impact gonococcal immune evasion. It is well established that LOS-sialylation and FH recruitment increase gonococcal resistance to killing by human serum [23]. Therefore, we hypothesized that exposure to VLY-extracted HeLa contents would potentiate *N. gonorrhoeae* resistance in a serum-killing assay. *N. gonorrhoeae* F62 was incubated in the presence of 4% normal human serum (NHS) and colony-enumeration was used to assess survival. As expected, there was a significant increase in gonococcal survival after exposure to the VLY-liberated HeLa cell contents compared to vector-only controls (Fig. 6A). This difference was not observed when *N. gonorrhoeae* F62Δ*lst* was tested (Fig. 6B), suggesting that the survival depended on LOS sialylation.

**Figure 6:**
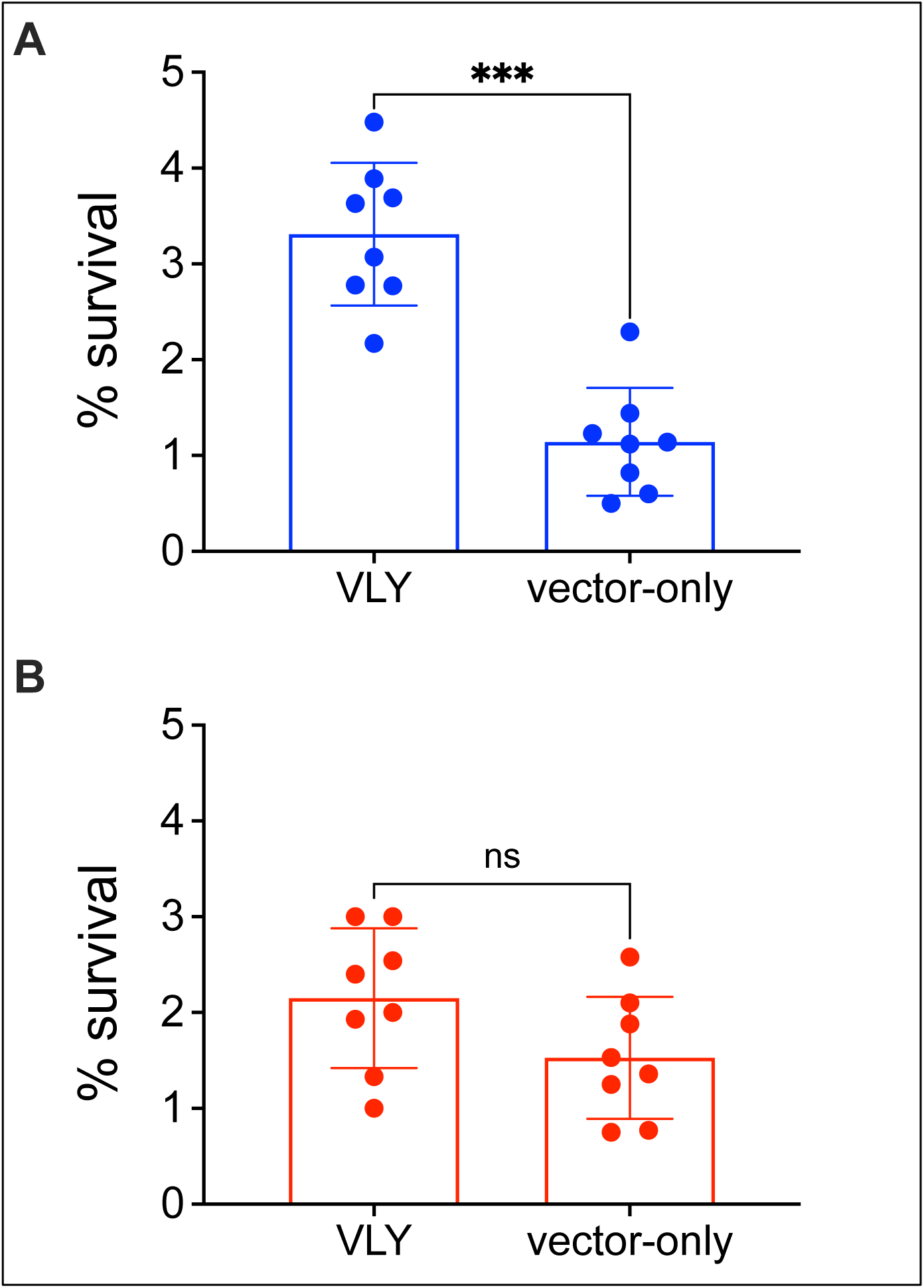
Exposure of *N. gonorrhoeae* to VLY-liberated HeLa contents increases sialylation-dependent survival in human serum. *N. gonorrhoeae* F62 WT and Δlst were incubated with the supernatants of HeLa cells that had been exposed to either 1 μg/mL vaginolysin or a parallel dilution of the vector-control preparation. (A & B) Survival of *N. gonorrhoeae* F62 WT (A) and Δlst (B) after 40-min incubation in 4% NHS. Percent survival determined by colony enumeration at 0-and 40-min. Significance determined by Mann-Whitney. ***p<0.001 ns = no significance

## Methods

### Bacterial Strains and Cell Lines

*N. gonorrhoeae* F62 and F62 Δlst were grown on 15% Chocolate Agar plates [G.C. agar base (Remel) + 15% laked sheep’s blood] and incubated for 17-19 hours at 37°C, 5% CO2. *Gardnerella* ATCC14019 was grown either on NYC III agar or liquid medium supplemented with 10% heat-inactivated horse serum. ClearColi^Ò^ BL21(DE3) cells (Biosearch Technologies) were cultured per manufacturer instructions.

### Protein Cloning and Expression

Truncated VLY [*vly*(tr)] without the signal sequence was constructed, following amplification of the *vly* gene from *G. vaginalis* ATCC14019 beginning at position A45, similar to a previously described method, but instead with a 6-His tag [24]. This construct was ligated into pET28a vector and transformed into ClearColi BL21(DE3) cells.

For purification of VLY, cells carrying *vly*(tr)*-*pET28a or the vector only (empty) plasmid were grown in Terrific Broth + kanamycin (50 µg/mL) shaking at 37°C to an OD600 of 0.4. Protein expression was induced with 1mM IPTG for 16 hours at 18°C before being pelleted and resuspended in Tris-HCL buffer (50 mM Tris HCl + 300 mM NaCl + 5 mM imidazole; pH 8.0), containing protease inhibitors (Roche EDTA free, 0.2 mg/mL lysozyme, and 160 µM PMSF. Cells were lysed in this buffer via sonication on ice 26 times, 15 sec each, at 10% amplitude. Clarified lysates were incubated with TALON Metal Affinity Resin cobalt beads (Takara), and beads were then washed with Tris-HCL (pH 8.0) buffers with increasing concentrations of imidazole. The bound proteins were eluted using Tris-HCL buffer (pH 8.0) with 300mM imidazole. Preparations were evaluated for purity using Coomassie SDS-PAGE, and protein concentrations were estimated using BCA Protein Assay Kit (Pierce). Prior to use in assays, protein preparations were buffer exchanged into 1X DPBS using PD Midi-Trap G-25 columns (Cytiva).

### Hemolysis

To wash and isolate intact red blood cells, 5 - 10 mL of washed single donor human RBCs in Alsever’s solution (Innovative Research Inc) were added to a 50ml conical tube, followed by 30 - 40 mL of cold DPBS (no calcium, no magnesium). The tube was gently inverted several times to mix RBCs with DPBS, before pelleting the RBCs at 375 x g, 4 °C, for 10 minutes. DPBS wash of the pellet was repeated until the supernatant was clear rather than pink, indicating the absence of lysed RBCs and free hemoglobin. After the final wash, RBCs were resuspended in cold DPBS to a calculated OD640 of 10, and 62.5 µL of this suspension was added to each well of a 96-well v-bottom plate. Dilutions of recombinant VLY (2.5 - 1000 ng/ml) or the vector-only control were made in DPBS. 62.5 µL of each dilution were added (in triplicate) to wells containing RBCs, to create a final protein concentration range of 1.25 - 500 ng/ml. The plate was incubated aerobically at 37°C, shaking at ∼300 rpm for 40 minutes followed by centrifugation at 375 x g, 4 °C, for 15 minutes. Supernatants were transferred to a clear, half-area polystyrene plate and heme release was determined by reading absorbance at 545 nm.

### Cytolysis (LDH-Release)

HeLa cells (purchased from ATCC) were seeded in a 96-well sterile, tissue culture treated plate and grown to ∼85 - 99% confluency in DMEM supplemented with 10% FBS. Prior to the assay, media was removed and cells were washed twice with DPBS (no calcium, no magnesium). Dilutions of recombinant VLY (0.25 - 6 µg/ml) or parallel dilutions of the vector-only control were made in DMEM without FBS or phenol red, and 150 µL of each dilution was added (in triplicate) to wells containing cells. The plate was incubated 1.5 hrs shaking gently at ∼175 rpm in 37°C, 5% CO2. High-lysis control wells had no VLY added, but received 7.5 µL of “lysis solution” from the Roche Cytotoxicity Detection Kit Plus 15 minutes before the end of incubation. Supernatants were removed and centrifuged at 200 x g for 5 minutes to pellet any lifted cells. LDH analysis was conducted on supernatants using a Roche Cytotoxicity Detection Kit Plus, per manufacturer instructions. Percent LDH release was determined by comparing sample values against high-lysis control.

### VLY treatment of HeLa cells to generate supernatants with intracellular contents

HeLa cells were grown to 85 - 99% confluence in a 100 mm sterile tissue culture treated petri dish. Prior to treatment, cells were washed twice with DPBS (no calcium, no magnesium). A 1 µg/mL solution of recombinant VLY (or equivalently diluted empty-vector preparation) was prepared in DMEM (no FBS, no phenol red), and 4 mL of this solution was added to washed cells. Plates were incubated at 37°C, 5% CO2, shaking at ∼175 rpm, for 1.5 hrs. Following incubation, supernatants were collected into conical tubes and centrifuged at 200 x g for 5 minutes to remove any lifted cells. Supernatants were then passed through a 0.22 um cut-off filter to remove any cell debris before being used in gonococcal sialylation experiments.

### Gonococcal Sialylation

Sterile-filtered HeLa cell supernatants (or exogenous CMP-Neu5Ac dilutions in DMEM) were combined 3:1 with 4X GC media, and 315 µL was added to wells of a 48-well tissue culture treated plate (Corning). An overnight plate of *N. gonorrhoeae* was scraped to make an OD600 0.5 suspension in assay medium (DMEM + 1X GC media) and 35 µL was added to each well for a starting OD600 of 0.05. The plate was then incubated for 3 hours at 37°C, 5% CO2, shaking at ∼300 rpm. After incubation, bacteria were collected and transferred to a 96-deepwell (2 mL) plate (Fisher Scientific) and washed three times with wash buffer (1 mL of DPBS (no calcium, no magnesium) + 0.2% BSA). In order to minimize any contaminating signal from sialylated glycoproteins in the wash buffer, IgG-free BSA (Sigma) was used. After the final wash, bacterial samples were resuspended in wash buffer and each sample was split into 2-3 groups for use in different downstream assays.

### Lectin Binding

To create lectin pre-conjugation solution, biotinylated lectins (MAL-I and ECA) were diluted 1:500 into staining buffer (DPBS (with calcium and magnesium) + 0.1% BSA) containing Cy5-streptavidin (1:1000). Lectins and Cy5-SA were incubated at room temperature for ∼ 1 hour, with end-over-end mixing.

Following gonococcal sialylation, washed bacteria were transferred to a 96-v-bottom plate, spun down, and resuspended in 50 µL pre-conjugated lectin solution. Bacteria were incubated, shaking at 37°C for ∼50 minutes, then fixed with 1% formalin. Lectin-binding was assessed via flow cytometry using the BD FACSCanto^TM^ II High Throughput Sampler option.

### Factor H Binding

Following sialylation, bacteria were transferred to a 96-well v-bottom plate, spun down, and resuspended in wash buffer containing either 0.1 mg/mL Fc3/FH*, or 0.1 mg/ml Fc3 alone [21]. The plate was incubated, shaking at 37°C for 1 - 1.5 hours before cells were spun down and resuspended in a 1:000 dilution of FITC-conjugated anti-human IgG in wash buffer. Secondary incubation was done shaking at room temperature for 30-40 minutes. Bacteria were then fixed in 1% formalin and analyzed by flow cytometry as described for lectin binding.

### Serum Resistance

Following sialylation, 10 µL of washed bacterial samples (∼6×10^5^ CFU) were transferred to a 96-well plate and exposed to 4% pooled normal human serum (NHS) (Complement Technologies) in wash buffer. From each well, 10 µL was immediately removed and used to perform serial dilution and colony enumeration (t=0). The 96-well plate was incubated at 37°C, 5% CO2, shaking at ∼300 rpm for 40 minutes. At the end of the incubation, 10 µL was again removed from each well and used for CFU enumeration (t=40). Percent survival was determined by dividing the CFU at t=40 by the CFU at t=0, and multiplying by 100.

## Discussion

In this study, we asked if *Gardnerella*, an abundant bacterial genus in BV, could enhance the virulence *of N. gonorrhoeae* by providing access to intracellular contents. Exposure of epithelial cells to the *Gardnerella* cytolysin, VLY, enhanced gonococcal acquisition of LOS sialic acid, engagement in factor H binding, and evasion of complement killing. A limitation of our study is that we were unable to directly observe and measure CMP-Neu5Ac release from the HeLa cells upon treatment with VLY. However, CMP-Neu5Ac is the activated nucleotide sugar required for the gonococcal sialyltransferase, Lst. Thus, the data strongly support a model in which VLY-induced membrane damage result in enhanced access of gonococcal Lst to CMP-Neu5Ac otherwise contained inside host cells (Fig. 7).

**Figure 7:**
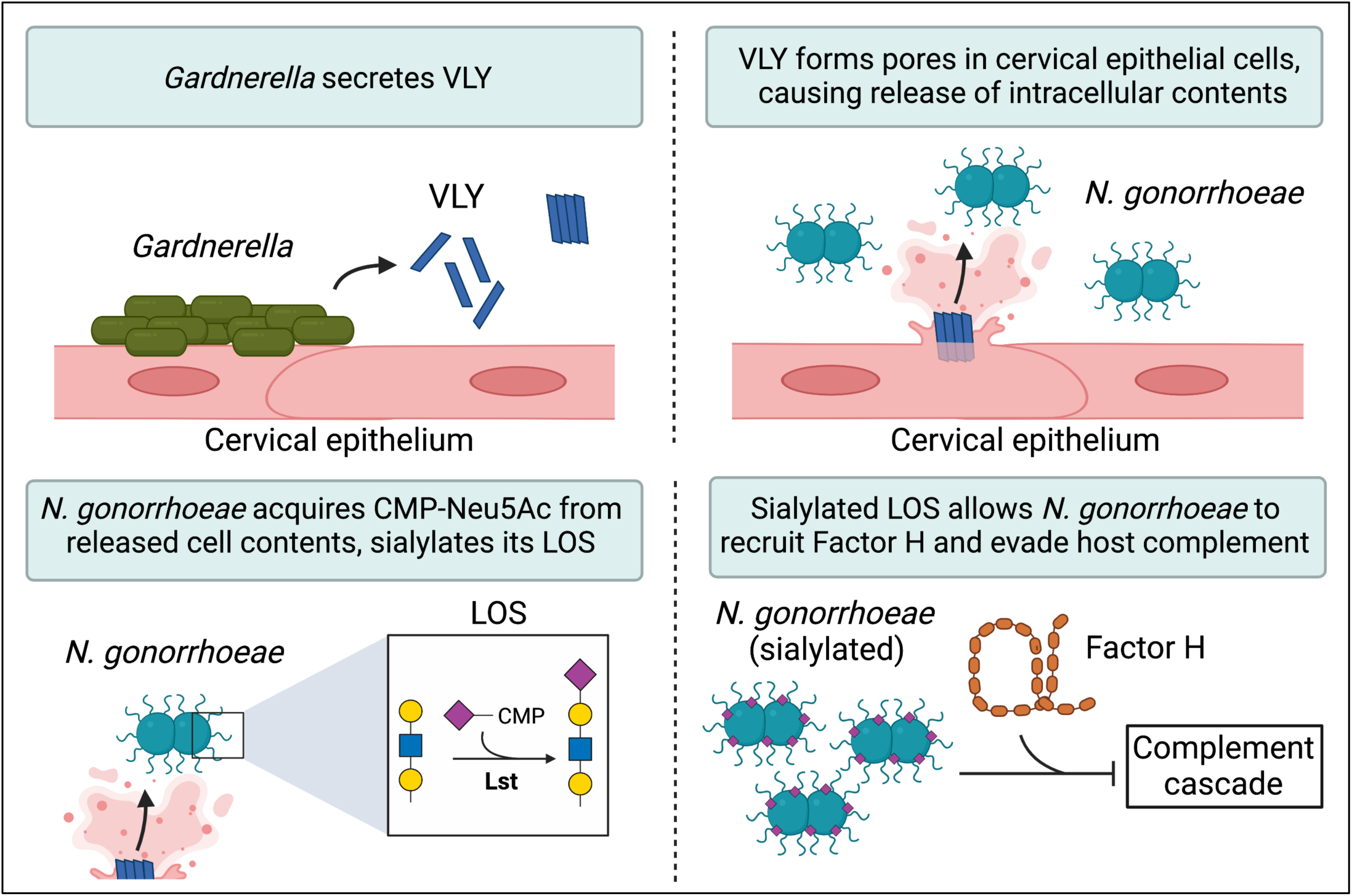
**Proposed model of** *Gardnerella* vaginolysin potentiating *N. gonorrhoeae* virulence. *Gardnerella* secretes VLY, which forms pores in cervical epithelial cells. Pore formation leads to a release of intracellular contents, including CMP-Neu5Ac. *N. gonorrhoeae* scavenges released CMP-Neu5Ac to sialylate its LOS, which enhances its ability to evade host immunity.

MAL-I recognizes the α2-3 sialylated *N*-acetylactosamine of LOS (Fig. 2A) and showed higher binding to the bacteria exposed to supernatants of VLY-liberated cervical epithelial cells compared to controls. Interestingly, MAL-I binding to F62Δ*lst* was still modestly higher on cells exposed to VLY-liberated supernatants compared to the vector controls (Fig. 3C-D). This could indicate the presence of molecules on the gonococcal surface with α-2,3-linked sialic acids independent of Lst activity. Alternative explanations include 1) the presence of mammalian glycans that were not fully removed by washing in the presence of VLY, 2) VLY-treatment releasing mammalian glycosyltransferases capable of sialylating gonococcal LOS [25], or 3) the bacteria produce a sialic acid-independent glycan epitope that binds to MAL-I (e.g. the lectin also binds 3-O-sulfated galactose[26]). Despite the uncertainty about the remaining MAL-I binding, the majority of the MAL-I binding was both VLY-and Lst-specific.

Molecular mimicry of host glycans is a well-characterized gonococcal virulence factor. LOS sialylation counters multiple host defense mechanisms including the classical and alternative pathways of complement, cationic antimicrobial peptides, and additionally dampens the inflammatory response by engaging Siglec receptors [20, 27-29]. In women, *N. gonorrhoeae* is believed to acquire CMP-Neu5Ac by invading epithelial cells. Our findings suggest that cell-damage driven by cytolysins present in BV may augment the ability of *N. gonorrhoeae* to scavenge this resource from the host, even during extracellular occupation of the niche.

A recent study showed that sialidases found in cervicovaginal secretions of women with BV can remove LOS sialic acids [30]. While de-sialylation of gonococcal LOS could render gonococci more susceptible to host defenses, it may confer other advantages. In fact, experimental evidence suggests that sialic acid removal may be required for asialoglycoprotein receptor binding and infection in men [31]. While Gardnerella sialidases can de-sialylate gonococci, our data show that their VLY may enhance the availability of CMP-Neu5Ac to promote LOS sialylation. Thus, the balance between sialidase and VLY activity may ultimately determine the net amount of LOS sialylation. Additionally, while almost all complete genomes of *Gardnerella* contain the gene for VLY, only a subset of strains encode sialidase genes and have sialidase activity in culture [32-34]. Future work should incorporate new experimental systems to investigate the interplay between sialidase and cytolytic activities in the gonococcal life cycle.

## AUTHOR CONTRIBUTIONS

SM helped conceive and develop the project, performed experiments, data analysis, wrote the first draft of the manuscript; SS performed experiments and data analysis, guided experiments, and participated in editing the manuscript; AV helped develop the project and participated in editing the manuscript, WG helped conceive and develop the project, guided experiments and data analysis and participated in editing the manuscript; SR helped develop the project and provided key reagents and funding for the project and participated in editing the manuscript; AL helped conceive the project, guided experiments and data analysis, provided funding for the project and participated in editing the manuscript.

## ACKNOWLEDGMENTS

We thank Dr. Keith L. Wycoff and Y Tran (Plant Biotechnology, Inc.) for providing Fc3/FH* (S2534)[21]. All schematics and illustrations were created with BioRender.com

## SUPPLEMENT

**Supplementary Figure 1:**
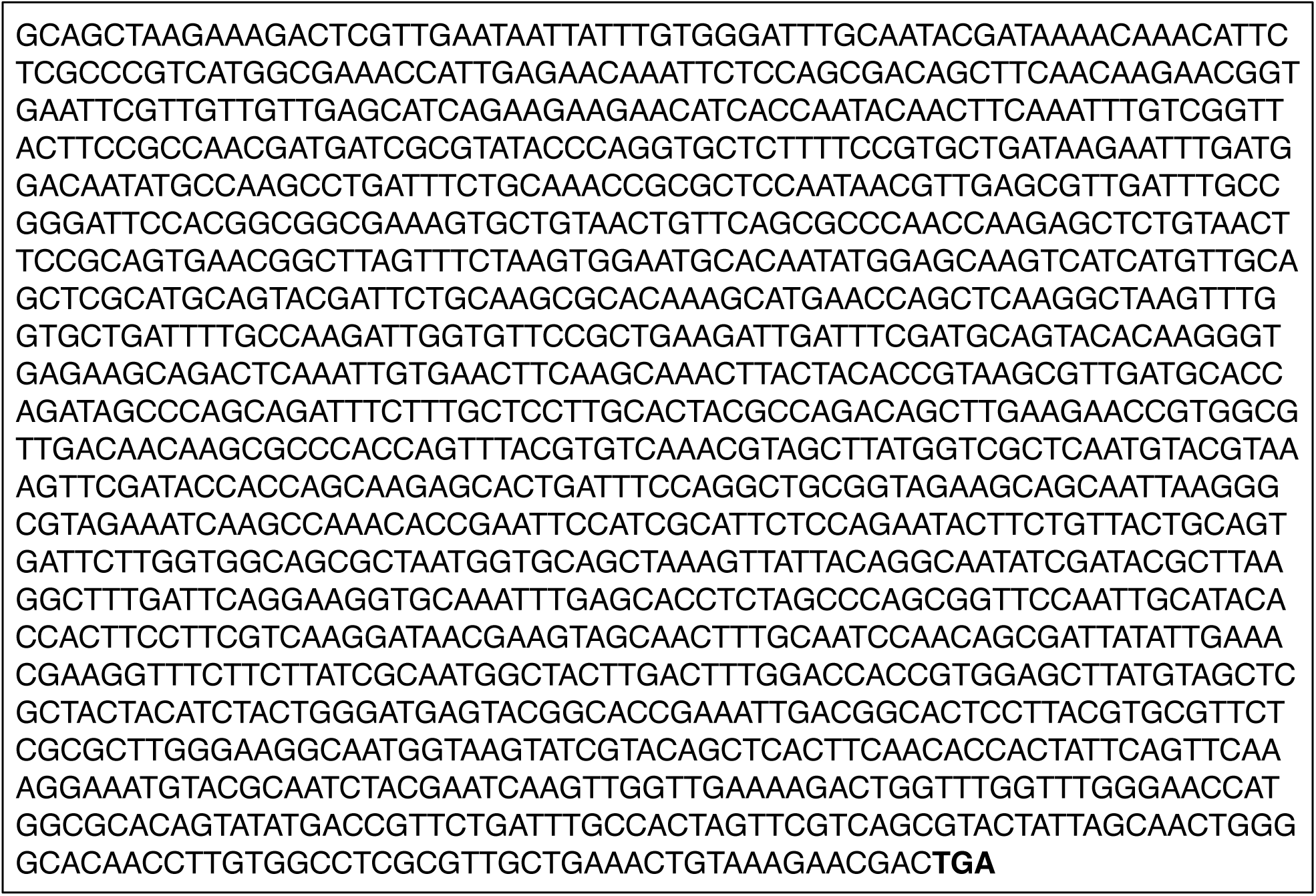
Nucleotide sequence of truncated VLY construct. Sequence source is the vly gene from *G. vaginalis* ATCC14109. Bolded nucleotides indicate stop codon.

**Supplementary Figure 2:**
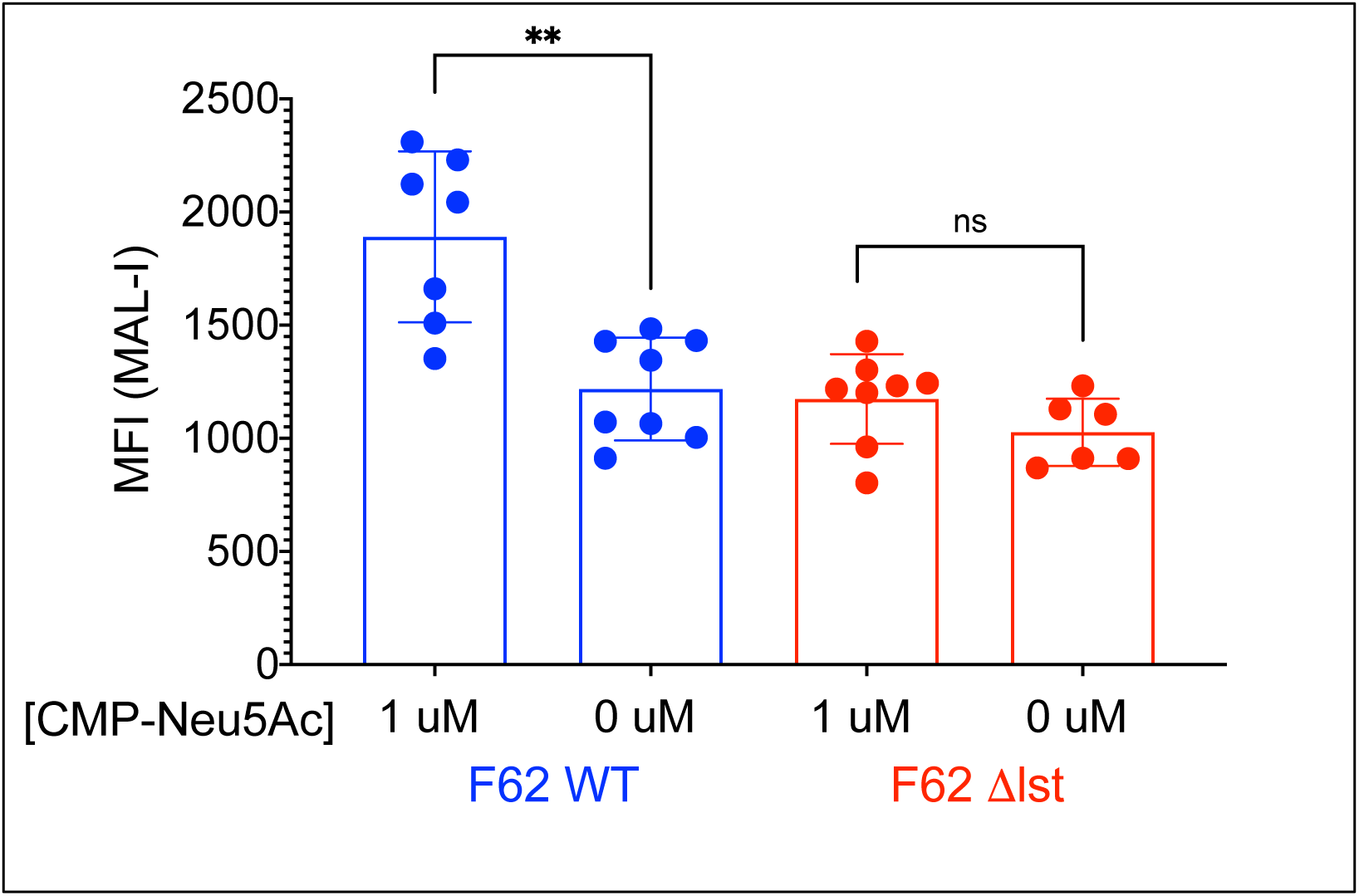
MAL-I binding requires Lst. (A) Binding of MAL-l I to *N. gonorrhoeae* F62 WT and Δlst following incubation with either 0 μM or 1 μM CMP-Neu5Ac. Presented as median fluorescence intensity (MFI). Data from two independent experiments, with three to four biological replicates per condition per experiment. Significance determined by Mann-Whitney. **p<0.01 ns = no significance

**Supplementary Figure 3:**
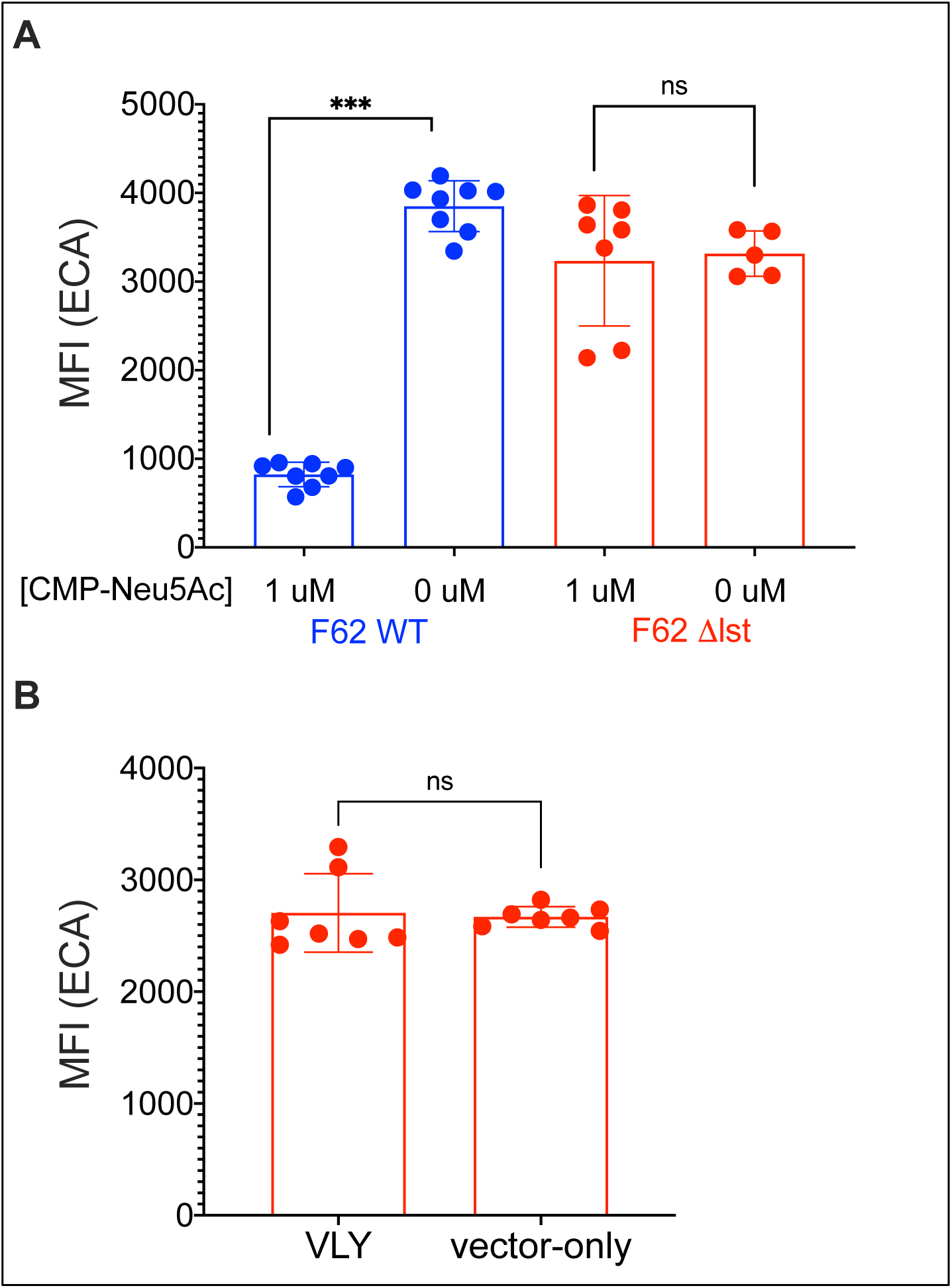
Decrease in ECA binding after gonococcal exposure VLY treated versus vector-treated HeLa cell supernatants is dependent on Lst. (A) Binding of ECA to *N. gonorrhoeae* F62 WT and Δlst following incubation with either 0 μM or 1 μM CMP-Neu5Ac. Presented as median fluorescence intensity (MFI). Data from two independent experiments, with two to four biological replicates per condition per experiment. (B) ECA binding after *N. gonorrhoeae* F62Δlst was incubated with the supernatants of HeLa cells that had been exposed to either 1 μg/mL vaginolysin or a parallel dilution of the pET28a vector-only control. ECA binding presented as median fluorescence intensity (MFI). Data is a combination of two independent experiments, with 3-4 biological replicates per experiment. (B & D) Significance determined by Mann-Whitney. ***p<0.001 ns = no significance

